# Genomic Similarity of Nucleotides in SARS CoronaVirus using K-Means Unsupervised Learning Algorithm

**DOI:** 10.1101/2020.10.12.336339

**Authors:** Jairaj Singh

## Abstract

The drastic increase in the number of coronaviruses discovered and coronavirus genomes being sequenced have given us a great opportunity to perform genomics and bioinformatics analysis on this family of viruses. Coronaviruses possess the largest genomes (26.4 to 31.7 kb) among all known RNA viruses, with G + C contents varying from 32% to 43%. Phylogenetically, three genera, Alphacoronavirus, Betacoronavirus and Gammacoronavirus, with Betacoronavirus consisting of subgroups A, B, C were known to exist but now a new genus D also exists,namely the Deltacoronavirus. In such a situation, it becomes highly important for efficient classification of all virus data so that it helps us in suitable planning,containment and treatment. The objective of this paper is to classify SARS corona-virus nucleotide sequences based on parameters such as **sequence length,percentage similarity between the sequence information,open and closed gaps in the sequence due to multiple mutations** and many others.By doing this,we will be able to predict accurately the similarity of **SARS CoV-2** virus with respect to other corona-viruses like the Wuhan corona-virus,the bat corona-virus and the pneumonia virus and would help us better understand about the **taxonomy** of the corona-virus family.

**SUMMARY:** In addition to the guidelines provided in the abstract above,the following points summarizes the article below:

- The article discusses an application of Machine Learning in the field of virology.
- It aims to classify the SARS CoV2 virus as per the already known sequences of the bat-coronavirus, the Wuhan Sea Food Market pneumonia virus and the Wuhan coronavirus.
- To solve and predict the similarity of the SARS CoV2 coronavirus w.r.t other viruses discussed above,**K-Means Unsupervised Learning** Algorithm has been chosen.
- The data-set used is **MN997409.1-4NY0T82X016-Alignment-HitTable.csv** found on **www.kaggle.com**.(Complete link shared in the references section).**[17]**
- The results have been validated by using a simple data-correlation technique namely **Spearman’s Rank Correlation Coeffecient**.
- I have also discussed my future work using **Deep Neural Nets** that can help predict new virus sequences and effectively find similarity if any with already discovered viruses.

## INTRODUCTION

Corona-viruses are single-stranded positive-sense RNA viruses that are known to contain one of the largest viral genomes, up to around **32 kbp**(base pair) in length**[1–5]**.In humans and birds, these viruses cause respiratory tract infections from mild range to lethal.With recent studies on the ever increasing genome sequences about the virus and the analysis made through prolonged research,the family of *Coronaviridae* can be classified into 4 distinct broad categories like the Alpha,Beta,Gamma and Delta versions of the corona-virus**[4,6,7,16]**. Alphacoronavirus and Betacoronavirus affect mammalian hosts, those in Gammacoronavirus and the recently defined Deltacoronavirus mainly infect avian species. As per the recent phylogenetic studies, it is seen that the coronaviruses have had a long history of evolution and mutation with frequent *zoonosis* leading to cross-species infections.Many of the corona-virus strains are known to be of the **bat origin** leading to a vast pool of recombination and mutation opportunities leading to cross-species transmission affecting mammals and humans notably**[4, 7, 8, 9, 10,16]**. One of the most unique characteristics of the corona-virus is its highly long genome base-pair sequence which can adapt any kind of new information it picks up in the process of mutation which makes the classification even more challenging. The most lethal **MERS and SARS** corona-viruses belong to the sub-genus *Sarbecovirus* and the *Marbecovirus* of Betacoronavirus genera**[9,11,12]**.Both of these result from the zoonotic transmission into humans leading to viral pneumonia,fever, breathing issues and other notable symptoms**[14,15]**.From the analyses of the whole genome to viral protein-based comparisons, it was found that the COVID-19 virus belongs to the lineage B (Sarbecovirus) of Betacoronavirus.From phylogenetic analysis of the *Rd-Rp* protein, spike proteins, and full genomes of the COVID-19 virus and other corona-viruses, it was found that the COVID-19 virus is most closely related to the *bat-like* corona-viruses**[12,16–20]**.There is also ongoing debate that the whether the COVID-19 virus developed as a recombination with previously identified bat and unknown corona-viruses **[16]** or arose independently as a new lineage to infect humans**[16]**.But with the identification of the *ACE-2* receptor in humans and its notable affinity with the corona-virus spike protein it has become quite evident that the COVID-19 virus is none other than a derivative of the bat corona-virus. The following model here utilizes an alignment-based approach using the very famous **K-Means Clustering** algorithm which relies on various annotations of the viral genes like sequence length,number of base-pairs present in the sequence and also the gaps or mutation affects on the information stored in the sequence.The benefits of using K-Means unsupervised learning algorithm lies in its easy of use in modelling complex data and high convergence rate resulting in accurate classifications.

## MATERIALS AND METHODS

The experiment performed uses a Machine Learning approach to classify the sequence of the *COVID-19* virus. I have used **K-Means Unsupervised Learning** algorithm primarily due to 2 factors:

- Due to the complexity of the data,comprising of various parameters which made it difficult to classify the virus into distinct labels.
- K-Means has a fast convergence rate and is easy to implement.

### Data Availability

The data-set used in the implementation can be found on Kaggle(Google’s ML platform)**(Link for Data Used[17])**.It is an accession-based data originally downloaded by performing a **BLAST** exhaustive search from NCBI(National Center for Biotechnology Information) database.The data has been described as per the following taxonomy.

- SARS-CoV-2(gene-bank)
- SARS-2019
- COVID-19 virus
- Wuhan corona virus
- Wuhan seafood market pneumonia virus.

Further,the columns in the table are further described as follows:

- MN997409.1 – – > SARS-CoV2-Accession Features
- MN997409.1.3 – – > Other SARS Accession Features
- 99.990 – – > percentage similarity between the virus strains.
- 29882 – – > the base pair alignment length of different virus strains.
- mismatches – – > mismatches in the sequence of SARS-CoV2 and other SARS accession features
- gap opens – – > open and closed gaps of the sequences,possibly due to mutation.
- q.start – – > the starting position of the SARS-CoV2 virus sequence
- q.end – – > the ending position of the SARS-CoV2 virus sequence
- s.start – – > the starting position of the other SARS-CoV2 virus sequence
- s.end – – > the ending position of the other SARS-CoV2 sequence. (see **Figure 1**)

**Figure 1.**
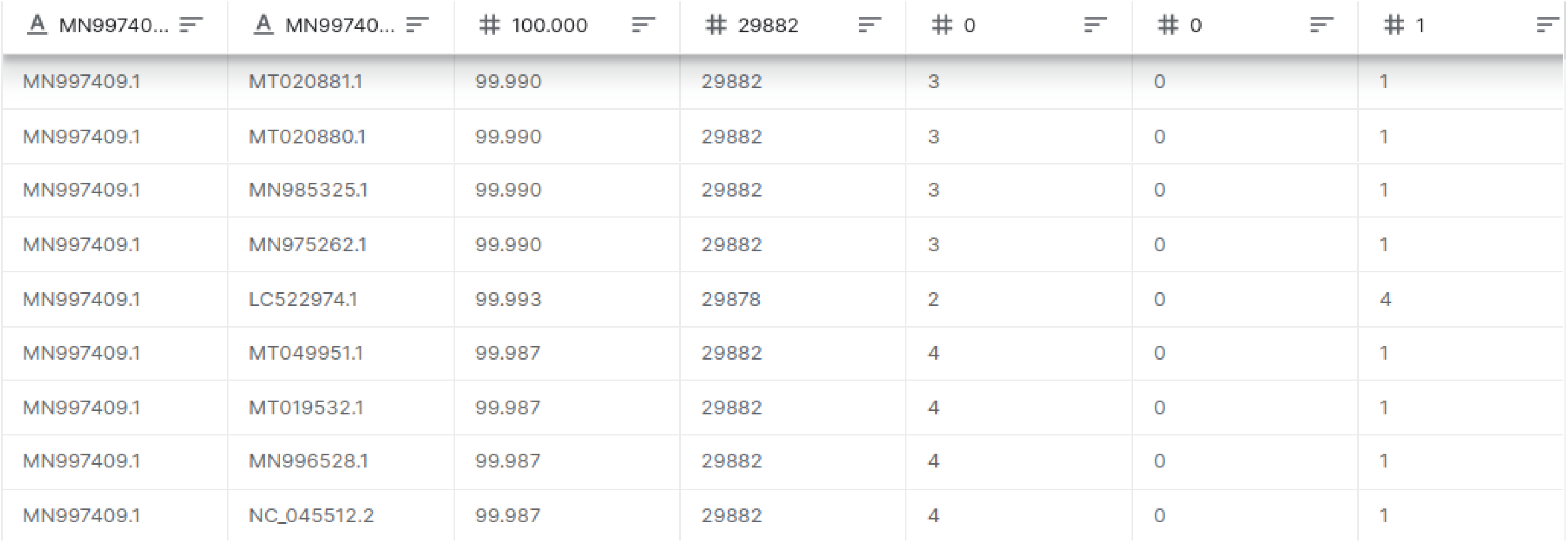
An overview of the data-table used in the experiment

### Statistical Analysis

The implementation of K-Means algorithm is based on the logic to find sequences that belong to the same cluster.If the cluster is found to be same then it implies that those sequences have large number of identical parameters.Since the data does not have any well defined labels or classes, therefore it has to be trained without human supervision,i.e.using an unsupervised learning paradigm.The working of K-Means goes by the selection of centroids.Centroids are first allotted randomly and then by running for a certain number of iterations the K-Means algorithm fixes the centroids and allocates the clusters.

But the question is how does the algorithm know where to stop ?.For,that it uses a performance-metric called as **inertia**,which is the **mean squared distance** between each instance and its allocated centroid.

The algorithm runs so as to always minimize the inertia.The K-Means can be mathematically represented as 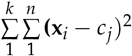 The *x*_*i*_ and *c*_*j*_ represent the instances and the allotted centroids respectively.There are two main kind of K-Means variants available namely *random* and *batch*.In random the centroids are allotted randomly and hence proceeds slowly whereas in batch K-Means the algorithm works much faster by avoiding unnecessary distance calculations.

### Optimal number of clusters

In order to find the optimal number of clusters for a K-Means algorithm, it is recommended to choose either of the following four methods:

- Elbow method (which uses the within cluster sums of squares)
- Average silhouette method
- Gap statistic method

In this experiment, I have used **Average silhouette** method to decide the optimal number of centers on this data-set.The Silhouette method measures the quality of a clustering and determines how well each point lies within its cluster.The main advantage that this method has it gives a visual representation of the number of clusters optimized as per data that makes it easy to implement K-Means effectively.

## RESULTS AND DISCUSSION

### Centroid Calculation

K-Means was implemented by exploiting the numeric values of the data such as the sequence length,percentage similarity, open and closed gaps. Since the data has 10 columns in total, K-Means was given a **random initialization** of centroids.This made the algorithm cruise through all the values gradually and thereby predicting the best centroids found after convergence from **10 iterations**. The metric got fixed at a value of **0.60536**(see approximately hence proving that algorithm reached the point of convergence. (See **Figure 2**). For the experiment,only the families with at least 100 sequences were considered.This is done so that maximum information is covered while performing the clustering process.If there are less than 100 sequences,it implies that the sequences lost their genetic information due to improper mutation or other *environmental conditions*.

**Figure 2.**
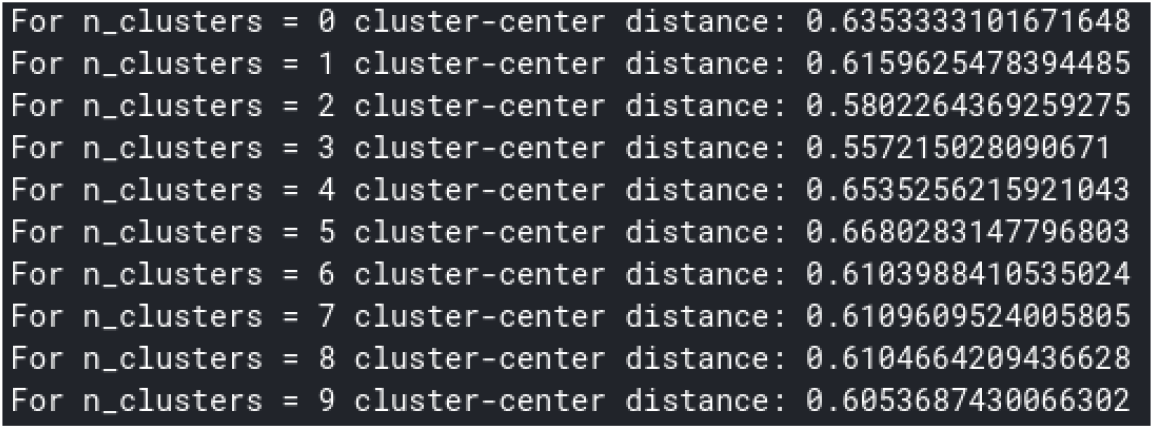
Computation of centroids through Silhouette method

### Cluster Formation

The cluster formation in K-Means is achieved through 2 steps:

- By using Silhouette distance to compute the cluster-center distance.
- By assigning appropriate labels,so that classification is easy to visualize and understand.

The algorithm as seen from above discussion ran for 10 iterations forming the best cluster in the last iteration.

The K-Means algorithm can also be accelerated using the **K-Means++** functionality in Python,where this model has been created. In the first iteration it can be seen that the clusters are oriented randomly as per the data, as the centroids have been randomly allotted.(see **Figure 4-Cluster Formation**)

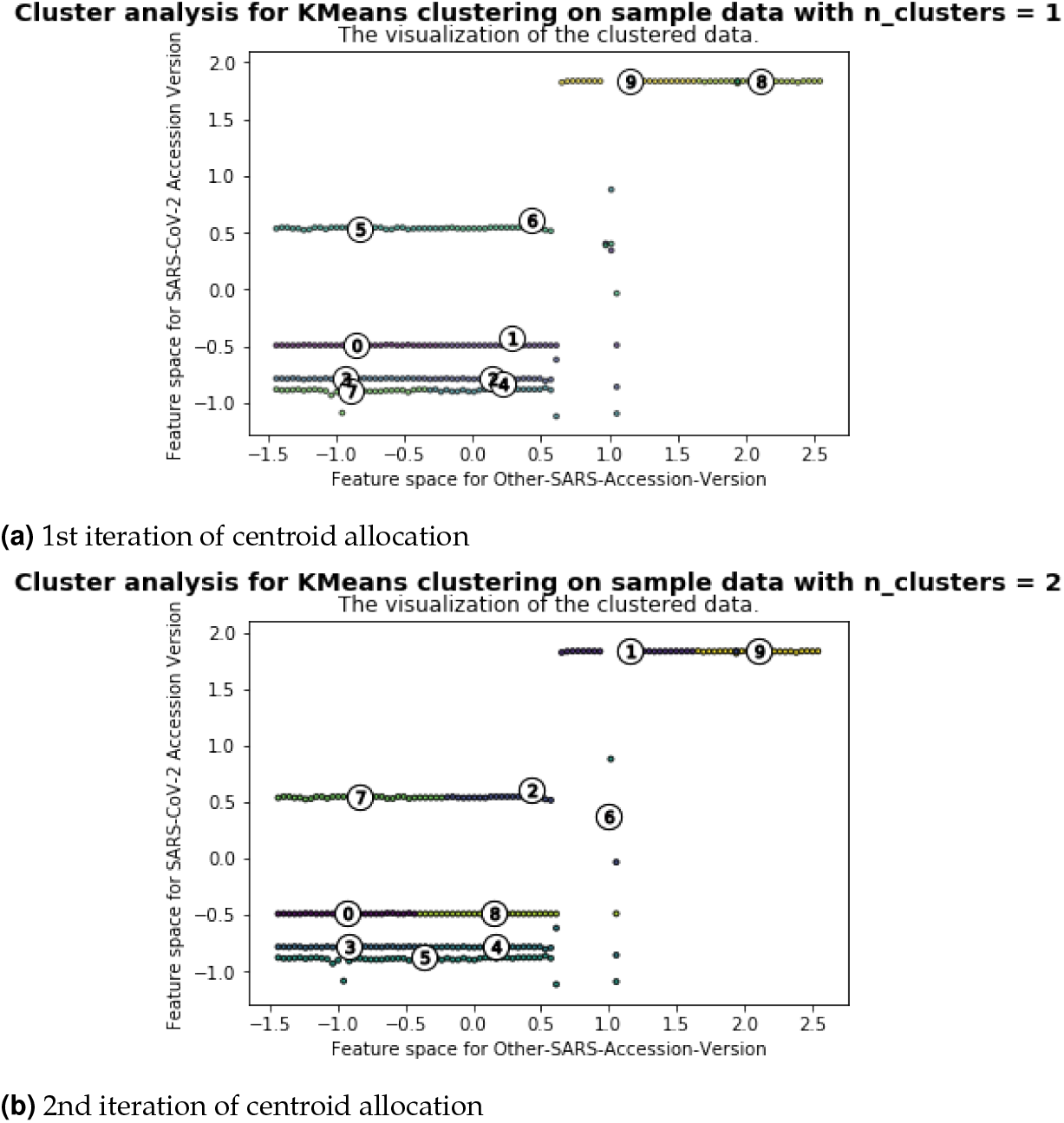

**Figure 4:**
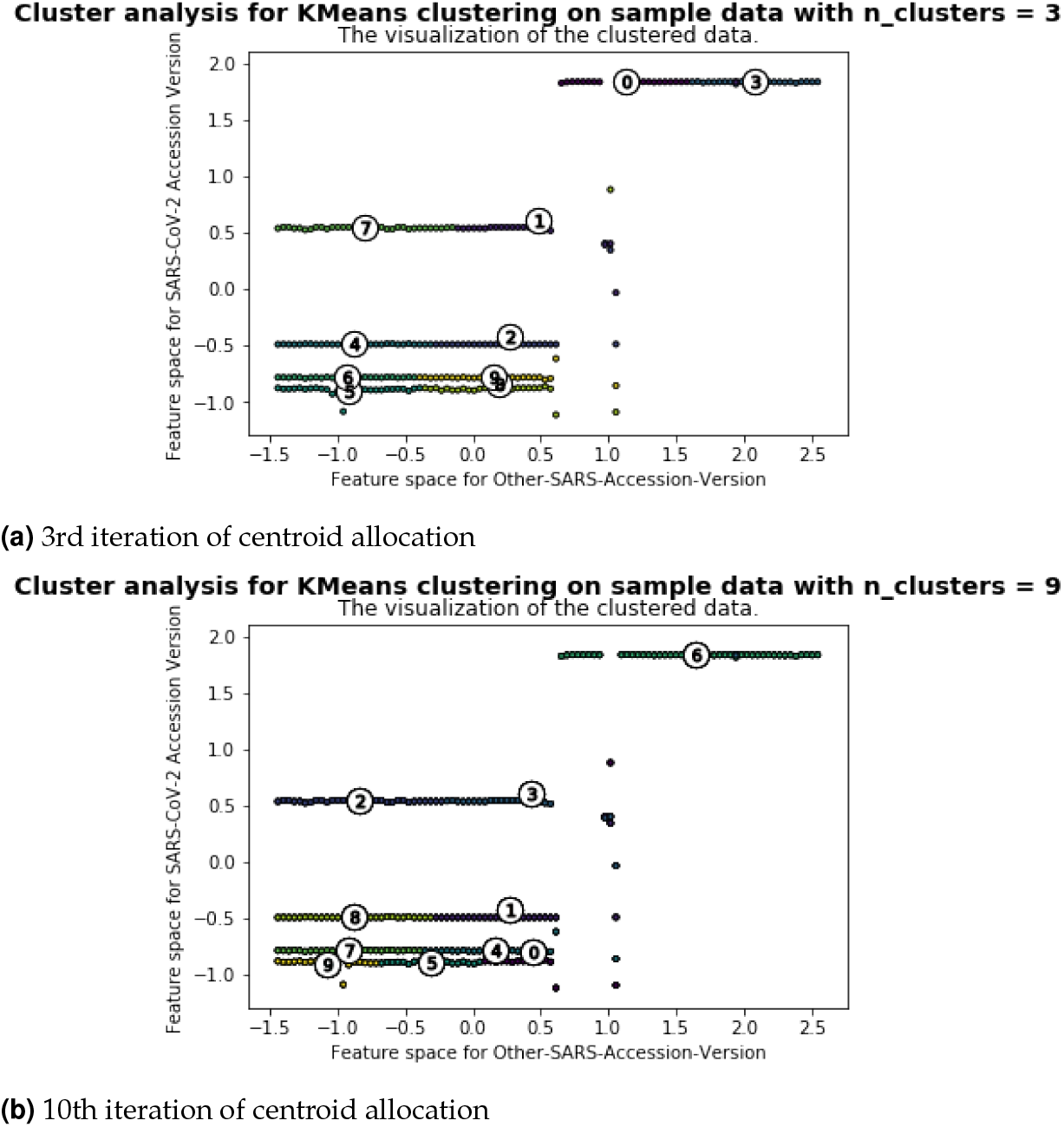
Cluster formation shown 1st,2nd,3rd and 10th iteration

In the second iteration it can be seen that the centroids got calculated and the clusters start aligning themselves accurately and it continues in this way all the way up to the 10th iteration.Here **only** the **first three iterations** and then the **final iteration** of cluster formation are shown.

Here the first three iterations and then the final 10th iteration are shown.What is to be noted here is that the clusters unlike in conventional methods are represented as circles,here it has been represented as **dots**.This has been done in order to output results the way a virus sequence appears naturally under a microscope to keep the visualization as natural as possible.

The diagram’s *x axis* represents the scaled feature space of the bat corona-virus,the Wuhan corona-virus and the Wuhan seafood market pneumonia virus while the *y axis* represent the current SARS-CoV-2 corona-virus feature space. Here the feature space is a scaled array of all instances described in the data.All these instances have numerical values and have been scaled within the range of −*1.5 to 2.0*.The scaling was automatically set by the program based on the *minimum and maximum limits* of the data used.

### Observation

The observation of the results show that clusters with centres numbered **9,5** and **7,4 and 0** show a high correlation with each other and thus belong to the SARS-CoV-2 sequence.The cluster with cluster numbered **6**(as per the diagram above in the 10th iteration) does not belong to the SARS-CoV2-sequence as its instances are **negatively correlated**.

The other cluster centers that lie in between share some information w.r.t the current COVID-19 virus but cannot be classified completely as a SARS-CoV2 sequence.They have a mix of other corona-virus strains such as the *Wuhan Sea Food Market Pneumonia virus* and the *Wuhan corona-virus*.

More notably,centers visible on the same strains signify that those cluster of viruses share exactly the same features and hence belong to the same class of the virus.

Thus we see that although the data-set was convoluted initially with no well defined labels,through this experiment we have been able to classify the virus reasonably into **labelled-clusters**.

### Validation

The results here also confirms w.r.t the heat-map shown above in the **results** section, the closeness of the COVID-19 virus with the sequences from the **Betacoronavirus genus** by a quantitative analysis based on the **Spearman’s rank correlation** used in the heat-map.(see **Figure 5-Heat-Map**)

**Figure 5:**
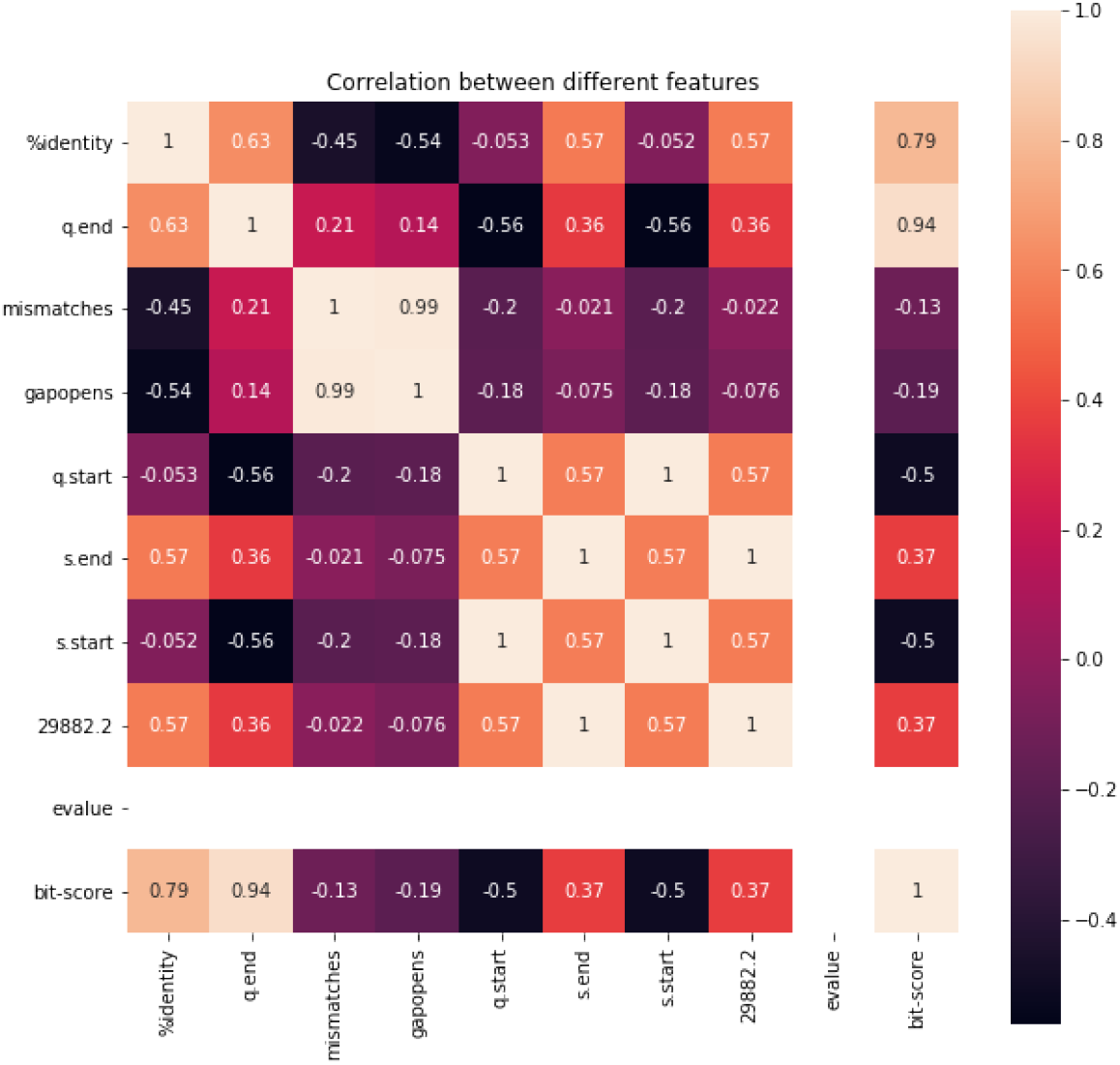
Spear-man’s Rank Correlation Heat-Map for result validation

Spear-man’s Rank Correlation is a metric describing similarity and dissimilarity between a pair of two ranked variables associated with each other.It is better than *Karl Pearson Correlation* as it does not require well defined parameters to find relationship between data. Also it works well on *categorical* data which remain fixed(*like age,length,mass*) throughout the experiment. In our data-set the correlation is between the target SARS-CoV2 virus and the other SARS variants found in either pneumonia virus and the Wuhan Seafood Market Virus.It is a categorical data set because it has **parameters** such as *length of bp*,*open-close-gaps*,*percentage similarity*.

Prior work elucidating the evolutionary history of the COVID-19 virus had suggested an origin from bats prior to zoonotic transmission **[12, 16]**. Most early cases of individuals got infected with the COVID-19 virus originating in the Huanan South China Seafood Market **[12–16]**. This proves that **Human-to-Human transmission** is confirmed, further highlighting the need for continued intervention.Still, the early COVID-19 virus genomes that have been sequenced and uploaded are over *99 percent* similar, suggesting these infections result from a recent cross-species event **[12,16]**.

## CONCLUSION

Although the classification could be achieved through K-Means,the model also has the following limitations:

1. The data used is an alignment-based data which means that it would require tremendous computational resources **(both space and time)** to process even a larger data set with more data features.
2. Better classification can be achieved using a group of algorithms such as **Linear SVM,Quadratic SVM and even a Polynomial SVM**.**[16]**

Apart from these, even **DBSCAN** algorithm can be used as there is no need to specify the clusters.DBSCAN can automatically find the right shaped clusters from the data,no matter how complex. However,this model makes use of an **alignment-based data** for obtaining a suitable classification of SARS-CoV-2 nucleotide sequences. But the model can be further improved by incorporating even larger data-sets particularly with the use of multiple classification algorithms. One more method, which I am currently working on is using Deep Neural Network. The clear advantage DNN has that it is able to extrapolate high level features from low level ones.The second advantage is that DNNs can be trained in either of the three fundamental models of machine learning names **Supervised Learning**,**Unsupervised Learning** and **Semisupervised learning**.

DNNs can also be integrated quite easily with **Natural Language Processing** tools by using *K-Mers algorithm(text based tokenizer)* to extract the best sequence tokens based on an optimized length.(**K=7** is the best in most cases). DNNs have a great role to play in the field of Artificial Intelligence as they hold the key to solve higher dimensional problems involving data complexity and providing diversified learning and higher efficiency. With the kind of AI we have today, computers would soon be able to help mankind achieve greater feats in the field of medical science.

